# Target2DeNovoDrugPropMax : a novel programmatic tool incorporating deep learning and *in silico* methods for automated *de novo* drug design for any target of interest

**DOI:** 10.1101/2020.12.11.421768

**Authors:** Rafal Madaj, Ben Geoffrey, Akhil Sanker, Pavan Preetham Valluri

**Affiliations:** University of Madras, Chepauk, Chennai 600 005, India; Centre of Molecular and Macromolecular Studies, Polish Academy of Sciences, Poland; SRM University, Tamil Nadu 603203, India; PSG College of Technology, Coimbatore, India

**Keywords:** drug design, automated drug discovery, in silico drug discovery

## Abstract

The past decade has seen a surge in the range of application data science, machine learning, deep learning, and AI methods to drug discovery. The presented work involves an assemblage of a variety of AI methods for drug discovery along with the incorporation of in silico techniques to provide a holistic tool for automated drug discovery. When drug candidates are required to be identified for a particular drug target of interest, the user is required to provide the tool target signatures in the form of an amino acid sequence or its corresponding nucleotide sequence. The tool collects data registered on PubChem required to perform an automated QSAR and with the validated QSAR model, prediction and drug lead generation are carried out. This protocol we call Target2Drug. This is followed by a protocol we call Target2DeNovoDrug wherein novel molecules with likely activity against the target are generated de novo using a generative LSTM model. It is often required in drug discovery that the generated molecules possess certain properties like drug-likeness, and therefore to optimize the generated de novo molecules toward the required drug-like property we use a deep learning model called DeepFMPO, and this protocol we call Target2DeNovoDrugPropMax. This is followed by the fast automated AutoDock-Vina based in silico modeling and profiling of the interaction of optimized drug leads and the drug target. This is followed by an automated execution of the Molecular Dynamics protocol that is also carried out for the complex identified with the best protein-ligand interaction from the AutoDock-Vina based virtual screening. The results are stored in the working folder of the user. The code is maintained, supported, and provide for use in the following GitHub repository

https://github.com/bengeof/Target2DeNovoDrugPropMax

## Introduction

The past decade has seen a surge in the range of application data science, machine learning, deep learning, and AI methods to drug discovery[1–12]. This greatly enhances the tool set already in offer from the computational era of drug discovery such as in silico modeling [13–18] and therefore a tool that incorporates the best of what both worlds of computational modeling and AI can offer to drug discovery can greatly help researchers in identifying new drug candidates. The presented work involves an assemblage of a variety of AI methods for drug discovery along with incorporation of in silico techniques to provide a holistic tool for automated drug discovery. When drug candidates are required to be identified for a particular drug target of interest, the user is required to provide the tool target signatures in the form of an amino acid sequence or its corresponding nucleotide sequence. The tool collects PubChem data registered against the target involving experimental activity value(pIC50) of compounds against the target and the molecular descriptors of the compound. With the collected data, the tool performs an automated QSAR and with the validated QSAR model, prediction and drug lead generation are carried out. This is followed by an automated in silico modeling and interaction profiling of the protein-ligand interaction involving the drug target and compounds generated as drug leads. This protocol we call Target2Drug. This is followed by a protocol we call Target2DeNovoDrug. We train a generative recurrent neural network called LSTM Chem with SMILES of compounds with activity against the target. The generative model generates novel molecules that are likely to be active against the target by virtue of the training dataset of the generative model being compounds that are active against the target. This is again followed by an automated fast virtual screen and in silico modeling protocol involving AutoDock-Vina and interaction profiling of the protein-ligand interaction involving the drug target and compounds generated de novo. Further in small drug molecule discovery, it is required that the generated molecules possess certain properties like drug-likeness, and therefore to optimize the generated molecules toward the required property we use a deep learning model called DeepFMPO [21], and this protocol we call Target2DeNovoDrugPropMax. This is followed by the fast automated AutoDock-Vina based in silico modeling and profiling of the interaction of optimized drug leads and the drug target. This is followed by an automated execution of a computationally more expensive Molecular Dynamics protocol that is also carried out for the complex identified with the best protein-ligand from the AutoDock-Vina based virtual screen. The results are stored in the working folder of the user. A demonstrated use of the tool has been shown with the target signatures of Tumor Necrosis Factor-Alpha, an important therapeutic target in the case of anti-inflammatory treatment. The future scope of the tool involves, running the tool on a High-Performance Cluster for all known target signatures to generate data that will be useful to drive AI and Big data-driven drug discovery

## Methods

The algorithmic workflow and architecture of Target2DeNovoDrugPropMax are shown Fig.1 below. The workflow begins with the collection of target signatures in the form of the amino acid sequence of the target or its corresponding nucleotide sequence. The BLAST protocol of Target2Drug identifies targets with identity with the user given target to collect data with linked PubChem data required to perform AutoQSAR(Quantitative Structure-Activity Relationship) based drug lead generation. The data involving the experimental activity (IC50) reported against the target of interest and the molecular descriptors of the active compounds is fetched from PubChem by the program to perform QSAR modeling and prediction. QSAR protocol followed by the program is as follows. It involves establishing a linear and non-linear statistical correlation between experimental activity(pIC50) and molecular descriptors of active compounds. A total of 8 molecular descriptors for each active compound against a target is downloaded from PubChem and a QSAR model is built with all possible combinations and selections of descriptors where nCr = 256, n = 8 and r = 2,3,4,5,6,7. The QSAR model with high statistical quality is chosen for prediction by choosing the QSAR model with R^2^ value being close to 1. With the validated QSAR model, the prediction is carried out on all PubChem compounds that are structurally associative to the active compounds and the top 50 compounds are chosen as drug leads. The predicted compounds satisfy the Lipinski’s drug likeness criteria. In this way, drug leads from the existing compound library for a given target of interest is generated. To generate drug leads that are novel compounds not present in present ligand libraries we follow the approach of Gupta et. al. [22] that uses a generative LSTM neural network to design drugs *de novo* for a particular target of interest. The SMILES of the PubChem compounds with reported activity against the target is used to train a generative LSTM neural network of Gupta, Anvita et al [22] to generate SMILES of new compounds that are not presently known and registered on PubChem. The newly generated compounds by the generative LSTM neural network are likely to be active against the target of interest as the neural network was trained with SMILES of compounds that are active against the target of interest. Further in small drug molecule discovery, it is required that the generated molecules possess certain properties like drug-likeness, and therefore to optimize the generated molecules toward the required property we use a deep learning model called DeepFMPO [21]. The de novo generated lead compounds generated from the generative LSTM model of Anvita et al are used as input for the DeepFMPO model. The DeepFMPO model uses an actor-critic reinforcement based learning policy to optimize the structures of the compounds towards drug-likeness. This is followed by an automtated in silico modeling protocol in used to further access the interaction of the he de novo generated drug like molecules and the drug target of interest. To access the interaction of the target and the lead compounds, the tool performs a fast and computationally cheap and efficient *In Silico* modeling and analysis of the protein-ligand interaction using AutoDock-Vina and stores the results in the working folder. Following this, a computationally more expensive molecular dynamics approach using GROMACS is used to study the protein-ligand interaction and complex formation for the protein-ligand complex associated with the lowest binding energy score from the AutoDock-Vina based virtual screening protocol. Our main program through the ‘runGromacs.sh’ bash script initiates an automated MD protocol which first identifies the complex with best protein-ligand interaction from the autodock-vina virtual screening using the ‘find_best_affinity.py’ and generates RMSD(Root Mean Square Deviation), RoG(Radius of Gyration), RMSF(Root Mean Square Fluctuation) plots which reveal the stability of the protein-ligand system based on the Molecular Dynamics simulation carried out for the protein-ligand system using the bash script ‘runGromacs.sh’. The ligand parameterization used in the automated MD protocol follows Bernardi et. al. [23] To demonstrate the use of the tool, a demonstrated use has been carried out with the target signatures of Tumor Necrosis Factor-Alpha, a drug target of importance for the anti-inflammatory disease [24]. The code is hosted, maintained, and supported at the GitHub repository given in the link below https://github.com/bengeof/Target2DeNovoDrugPropMax

**Fig. 1.**
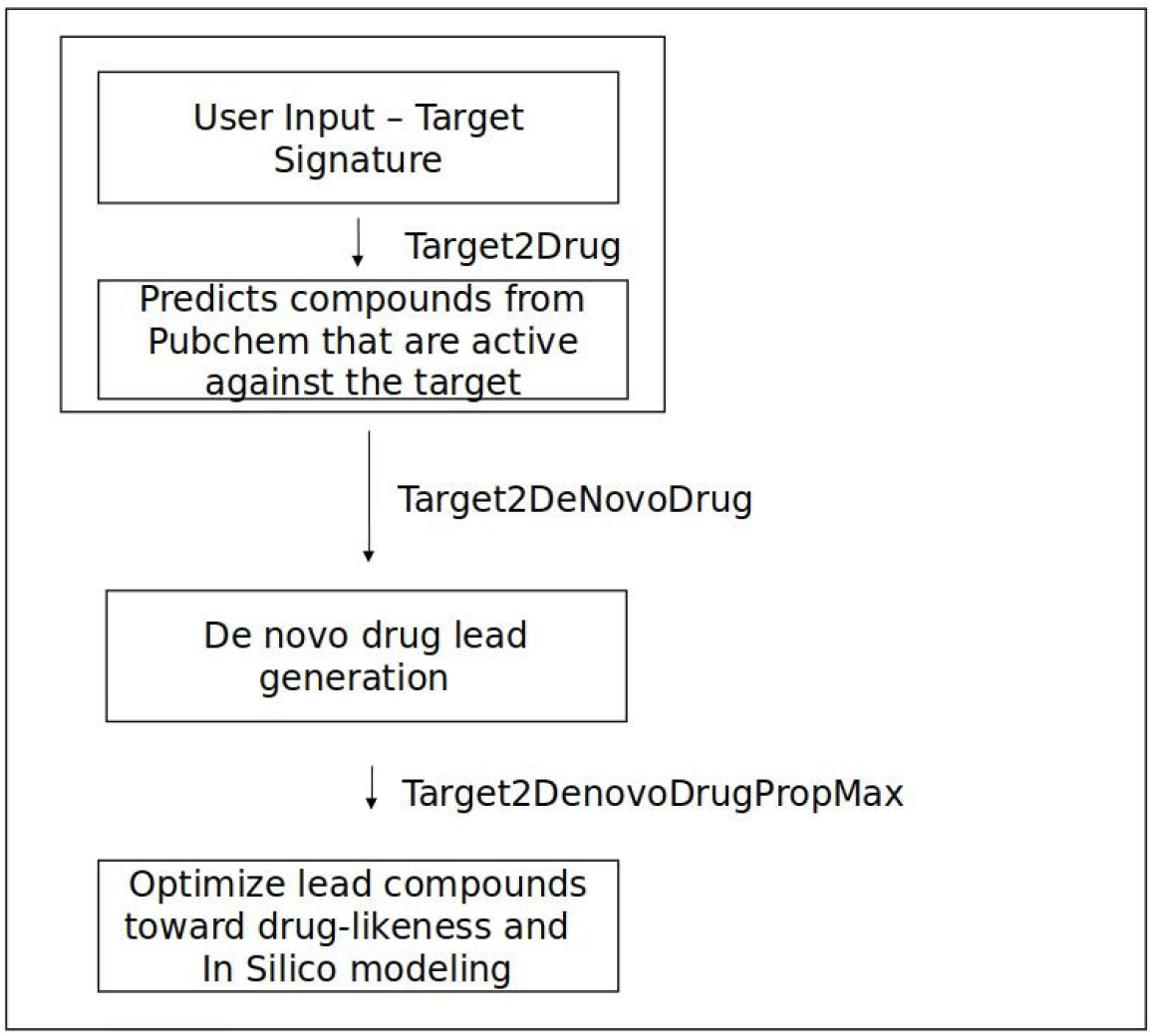
Block diagrammtic representation of the algorithm of the tool

## Results and Discussion

The target signature of Tumour Necrosis Factor Alpha was given as input to the tool. The target signature can be identified with NCBI protein ID P01375. The Target2Drug protocol identified data linked with the target on PubChem required to perform AutoQSAR based drug lead generation. The SMILES of the compounds generated as drug leads were used to train the generative LSTM neural network which was used to generate new compounds currently not known and registered on PubChem but are likely to be active against the target. 500 novel SMILES were generated out of which 182 were chemically valid SMILES. The newly Generated SMILES is shown in Fig.2. A computationally cheap and efficient *In Silico* modeling was carried out to model the interaction of the target and PubChem compounds generated as drug leads by the Target2Drug protocol and d*e novo* generated SMILES by the deep learning-based *de novo* drug design protocol.

**Fig. 2.**
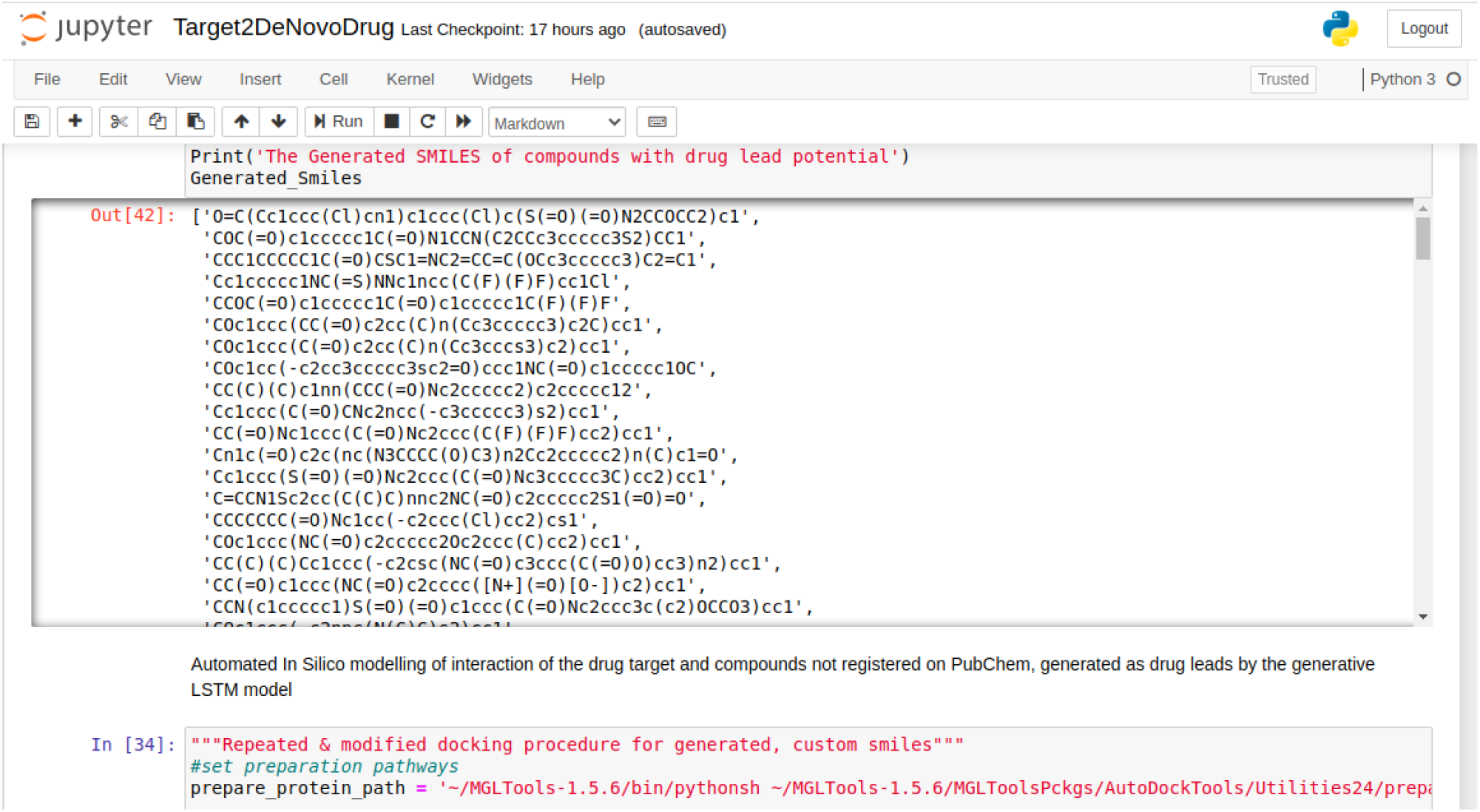
*de novo* generated SMILES

The results of the *In Silico* modelling is given the link below https://drive.google.com/drive/folders/1FFXQWPczR5hQfbruTF4YTb7K89wXcY9l?usp=sharing

A selection of drug lead compounds with Binding Affinity lower than −8 kcal/mol is tabulated and the pymol visualization of the target-compound interaction is shown in Fig.3a, b, c, d, e, 4a, 4b, 4c, 4d & 4e. The binding affinity of the target and PubChem compounds generated as drug leads is tabulated in Table 1 and the *de novo* generated compounds as drug leads are shown in Table 2 and drug like optimized lead compounds are shown in Table 3.

**Fig.3a.**
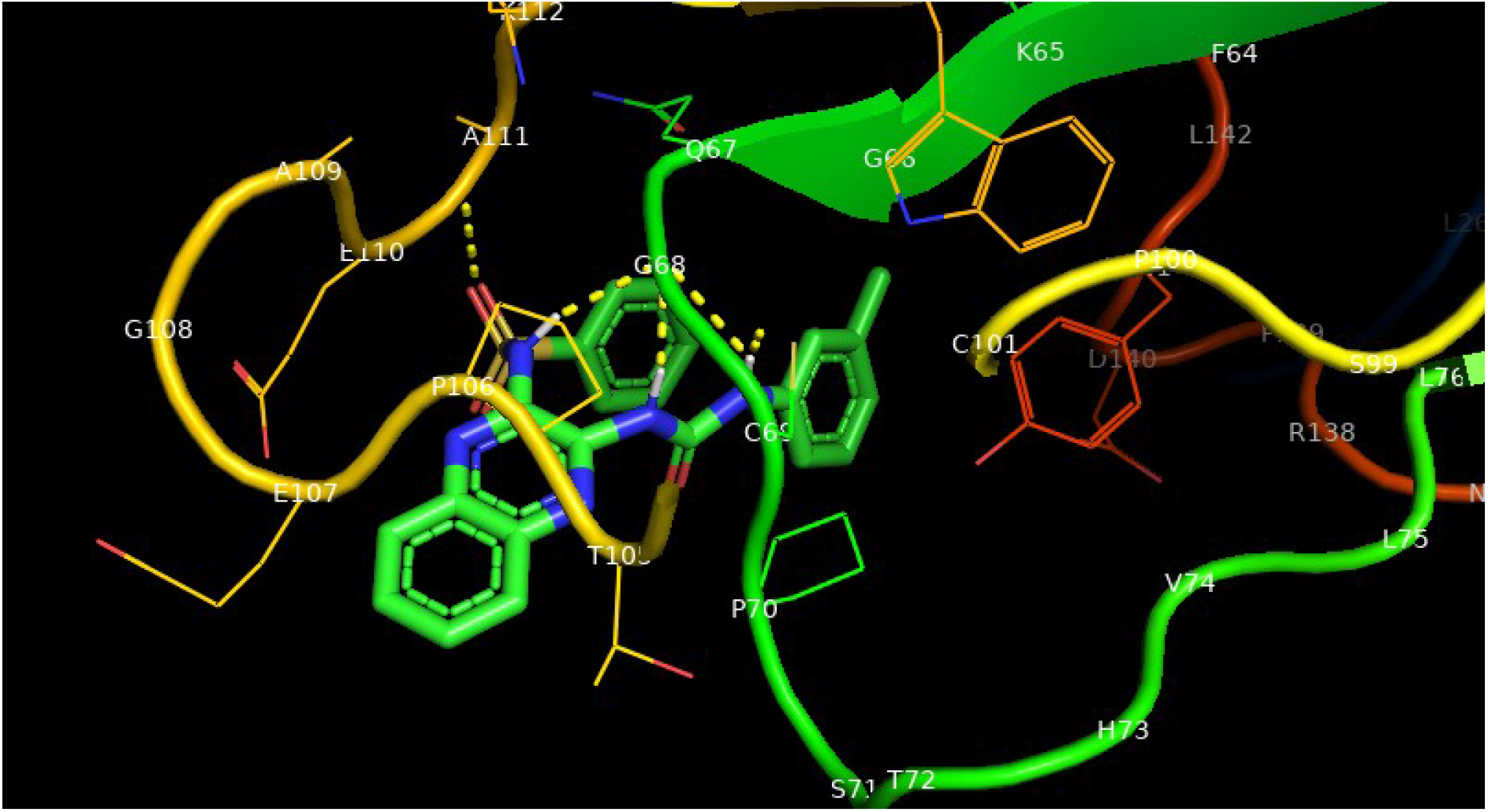
Target-compound interaction of the *de novo* generated SMILES 1-SMI

**Fig.3b.**
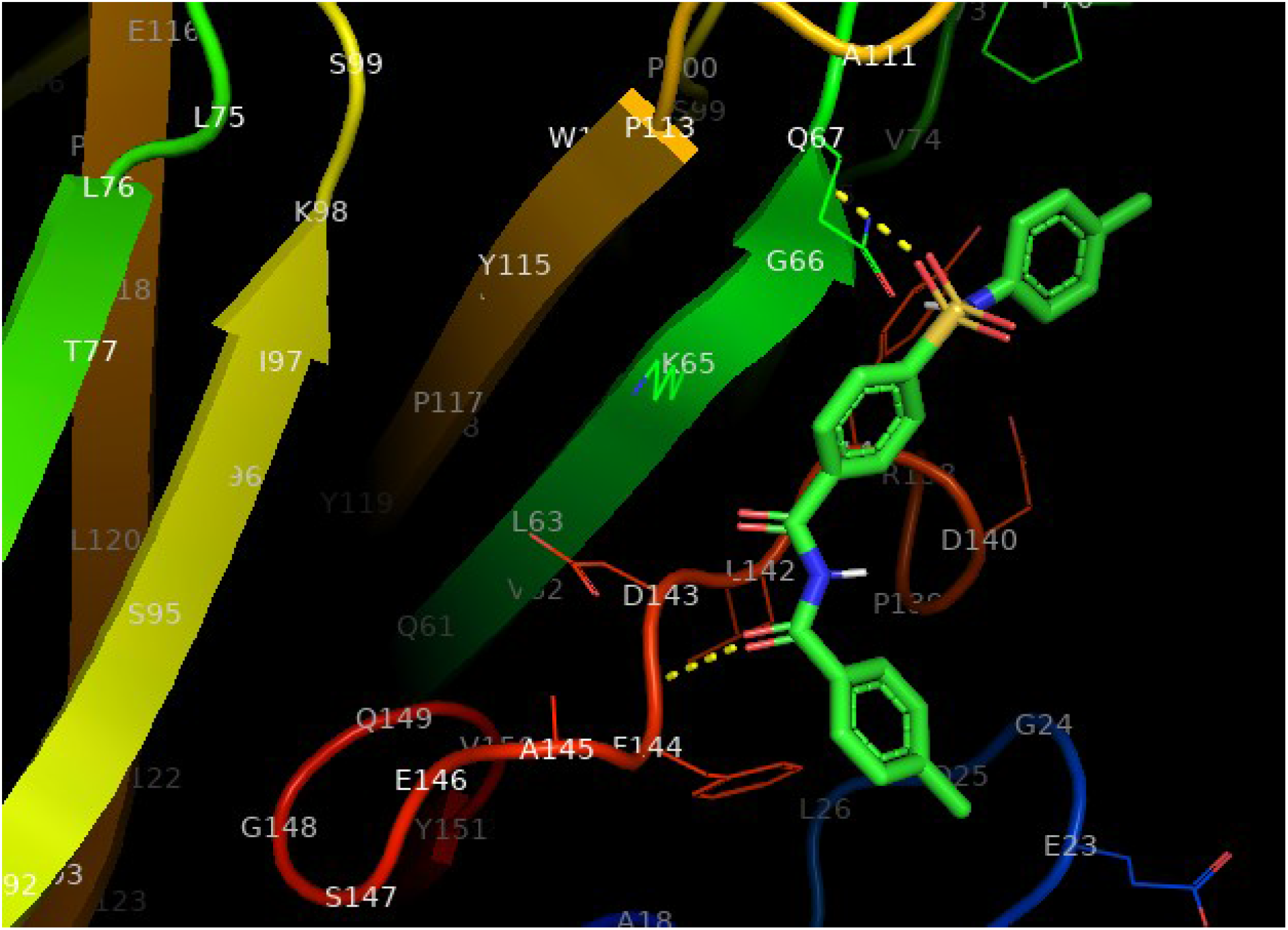
Target-compound interaction of the *de novo* generated SMILES 2-SMI

**Fig.3c.**
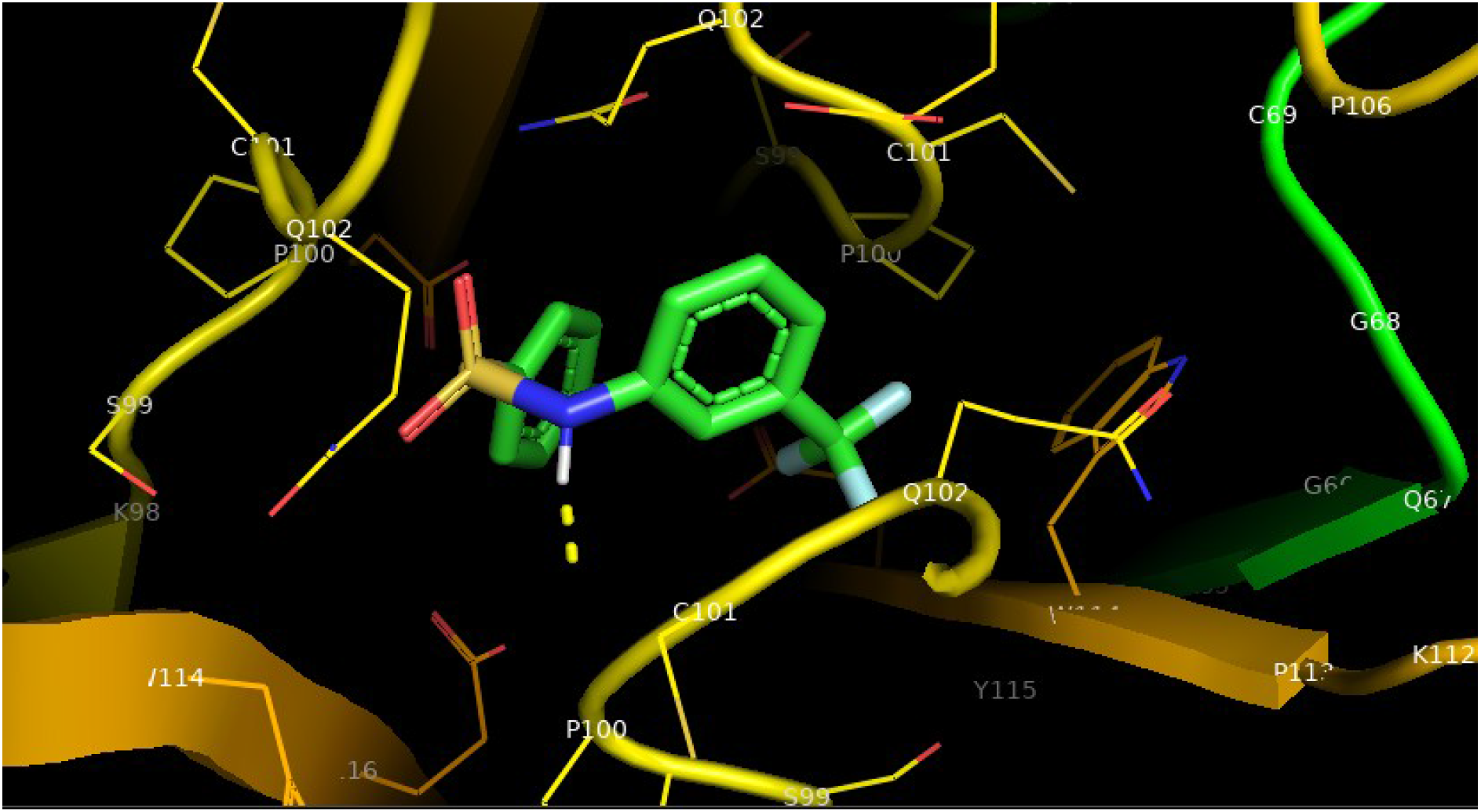
Target-compound interaction of the *de novo* generated SMILES 3-SMI

**Fig.3d.**
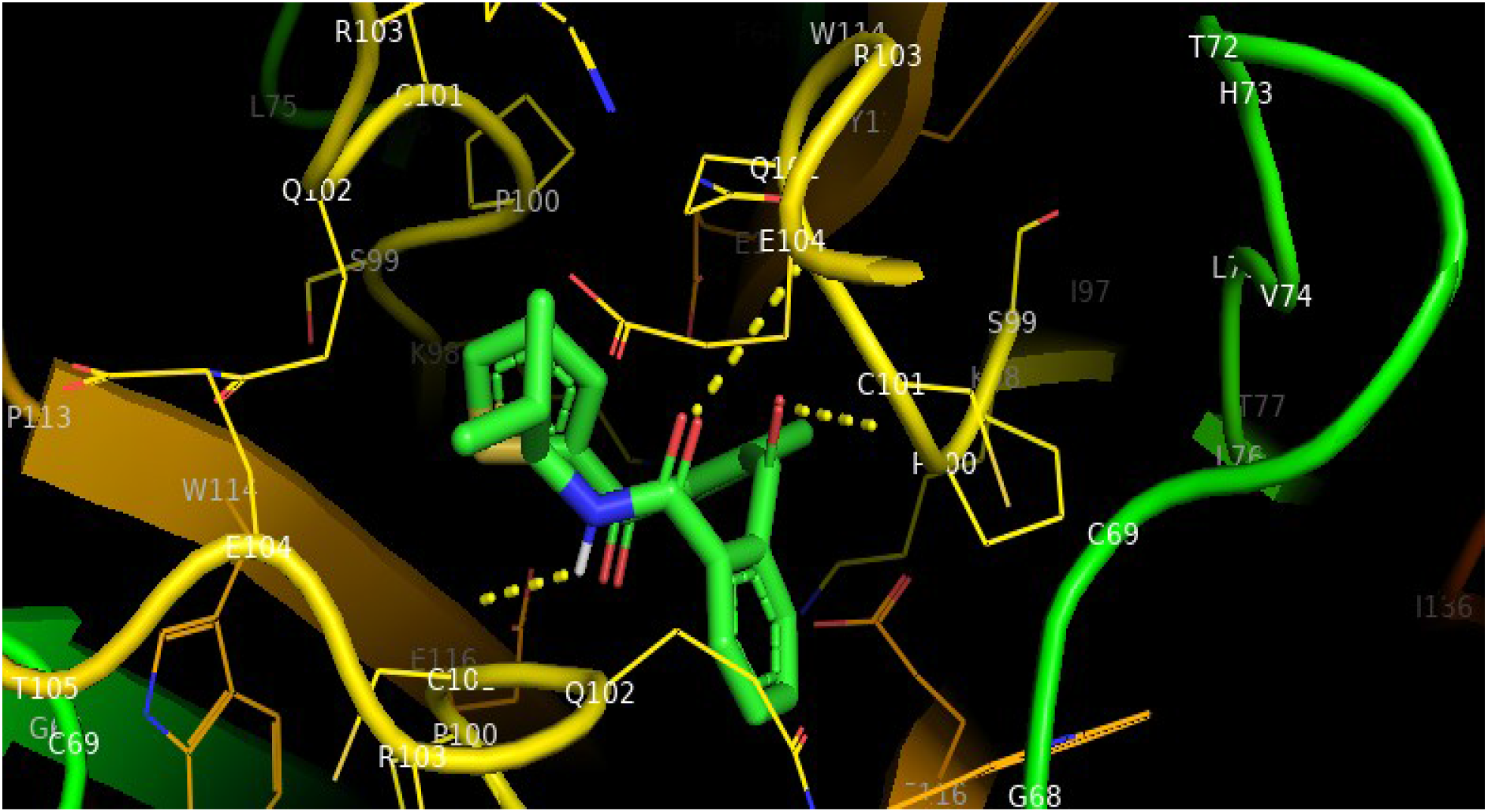
Target-compound interaction of the *de novo* generated SMILES 4-SMI

**Fig.3e.**
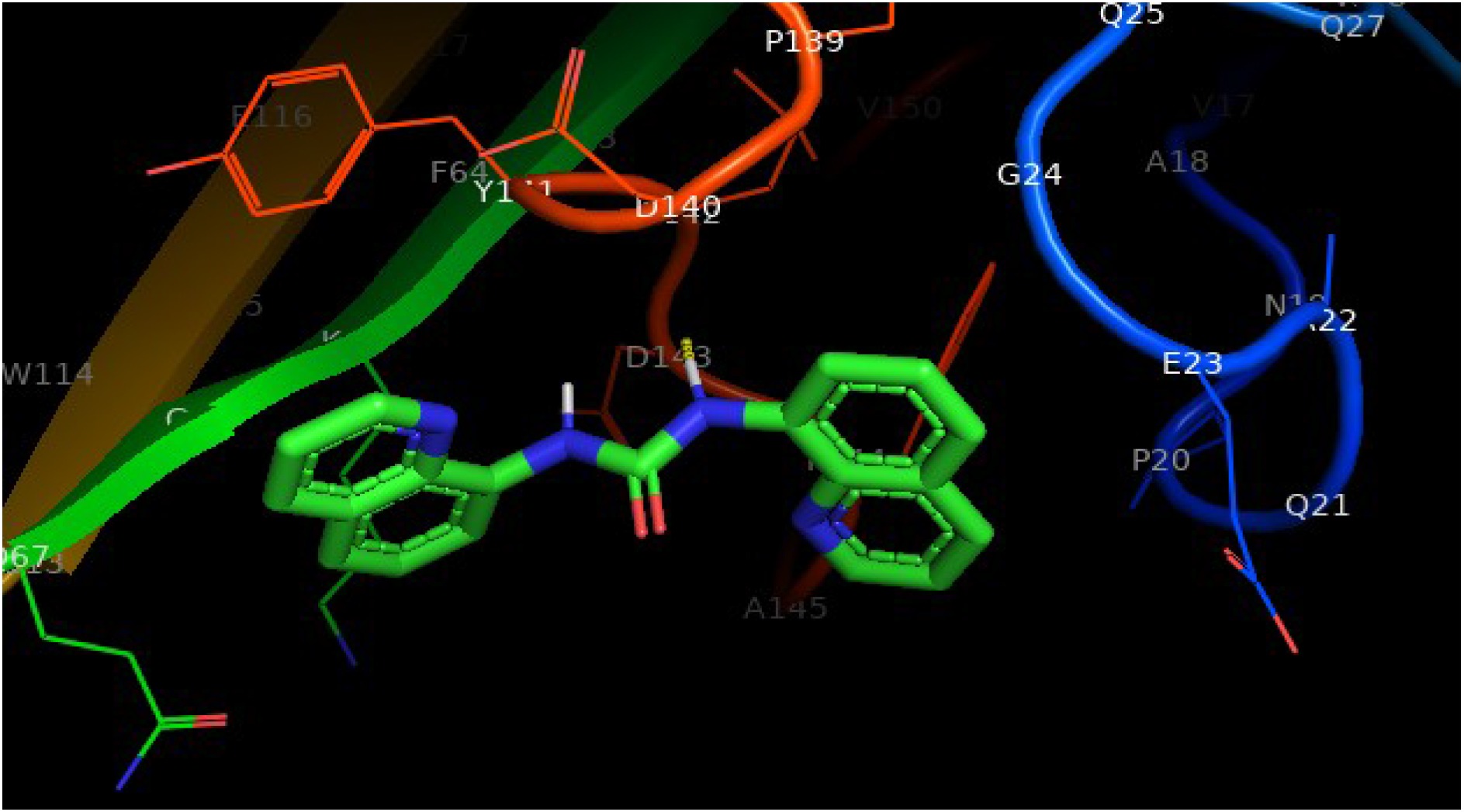
Target-compound interaction of the *de novo* generated SMILES 5-SMI

**Fig.4a.**
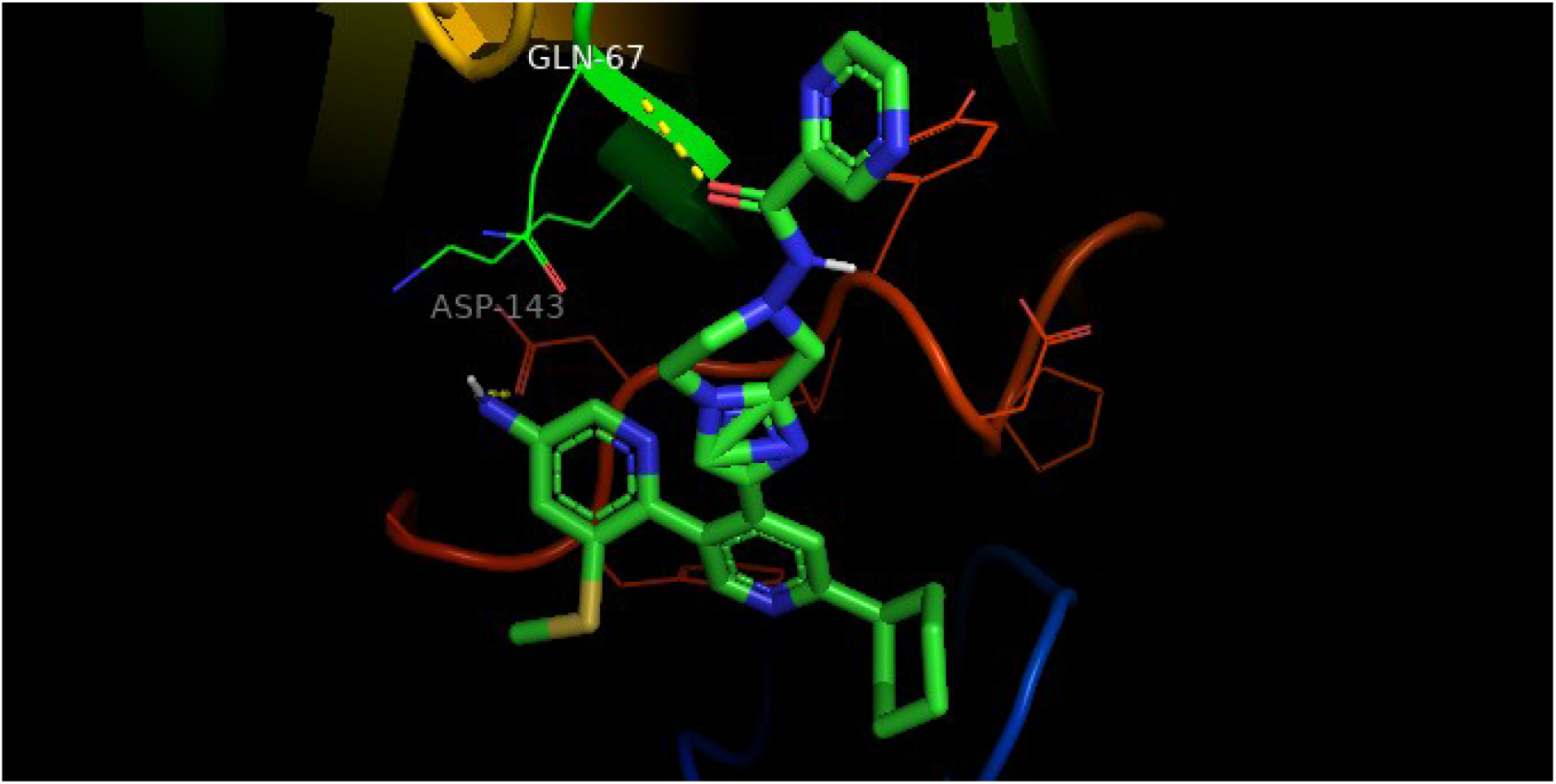
Target-compound interaction of the *de novo* generated SMILES 1-OPT-SMI

**Fig.4b.**
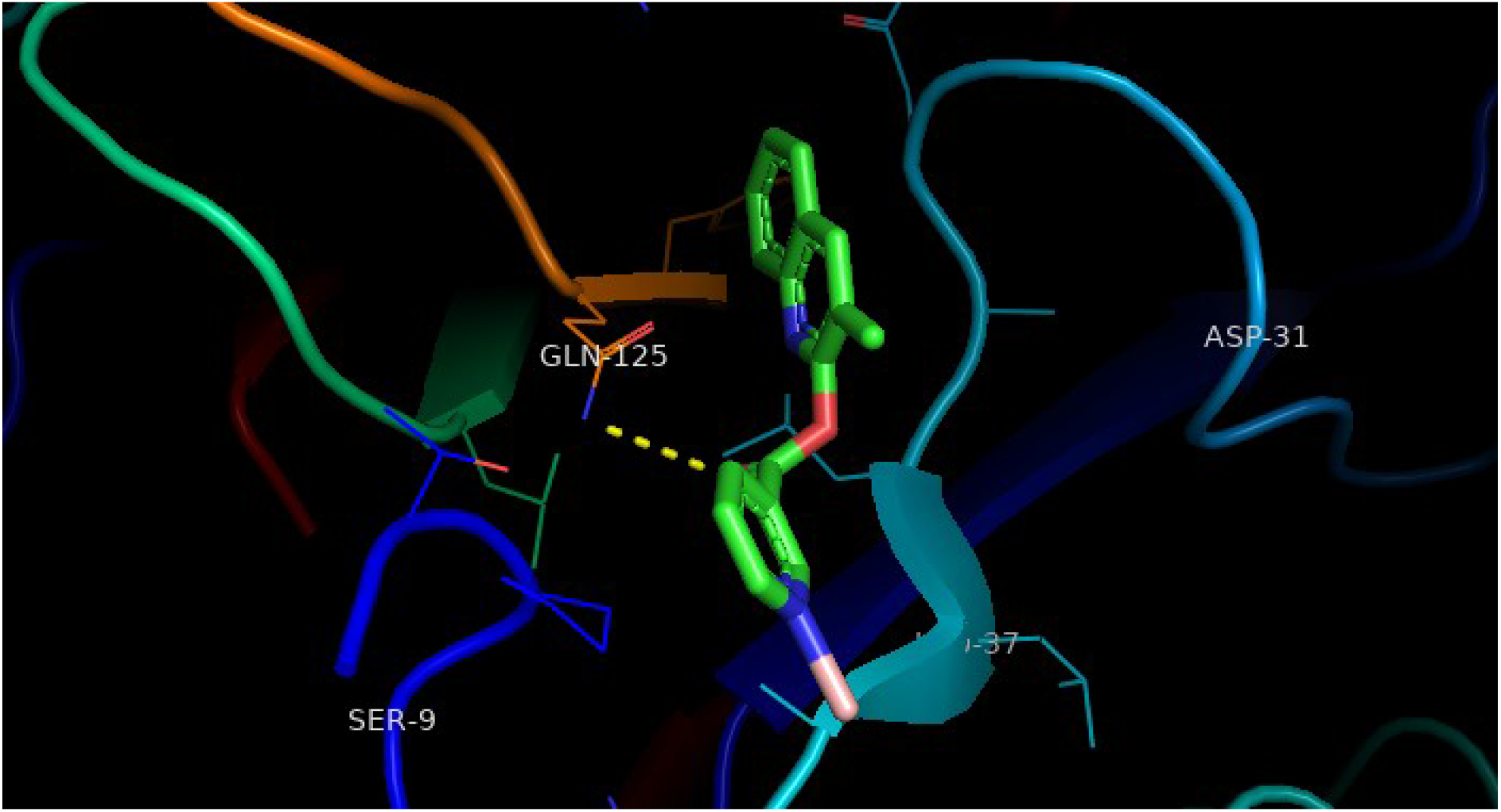
Target-compound interaction of the *de novo* generated SMILES 2-OPT-SMI

**Fig.4c.**
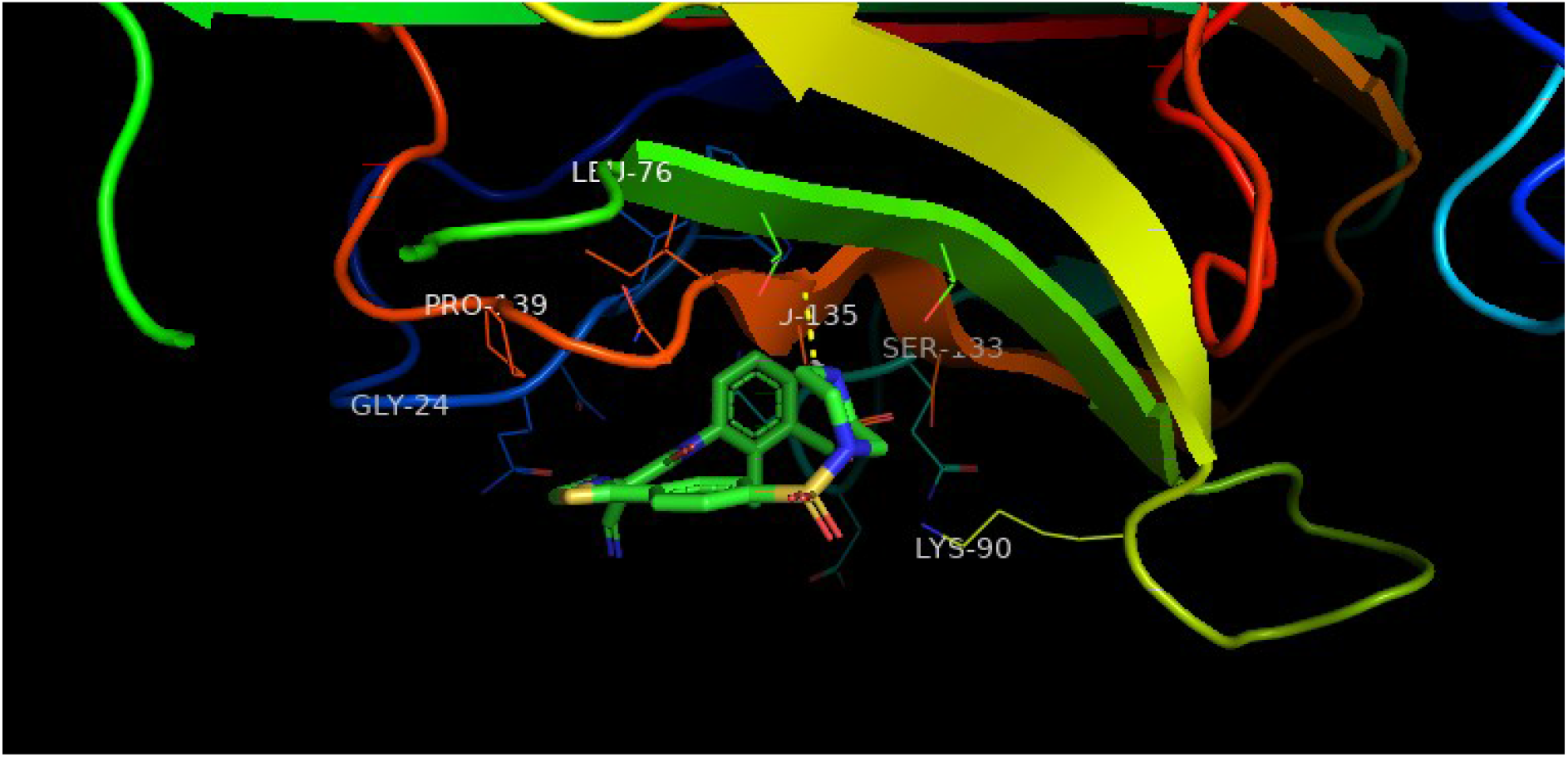
Target-compound interaction of the *de novo* generated SMILES 3-OPT-SMI

**Fig.4d.**
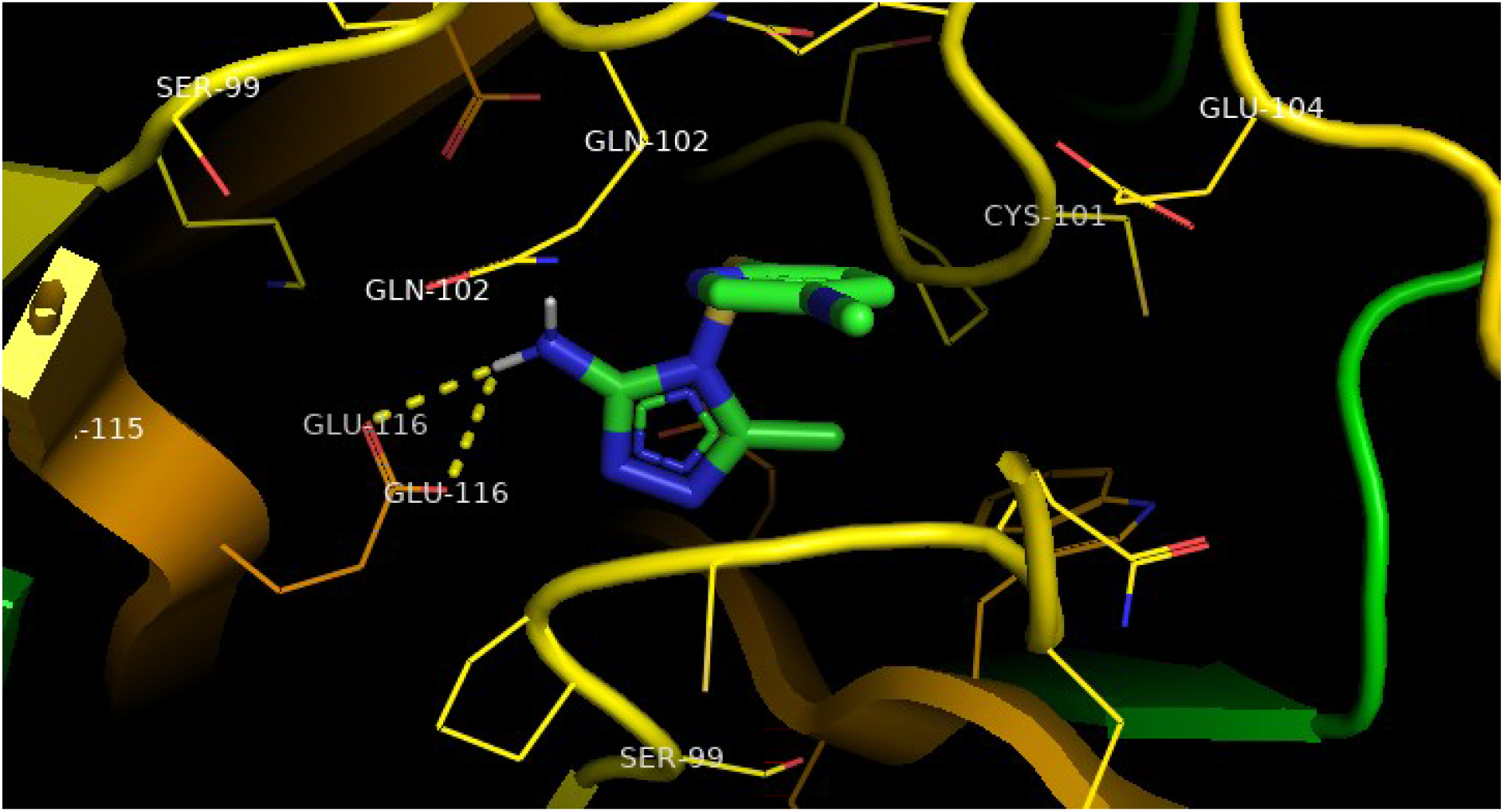
Target-compound interaction of the *de novo* generated SMILES 4-OPT-SMI

**Fig.4e.**
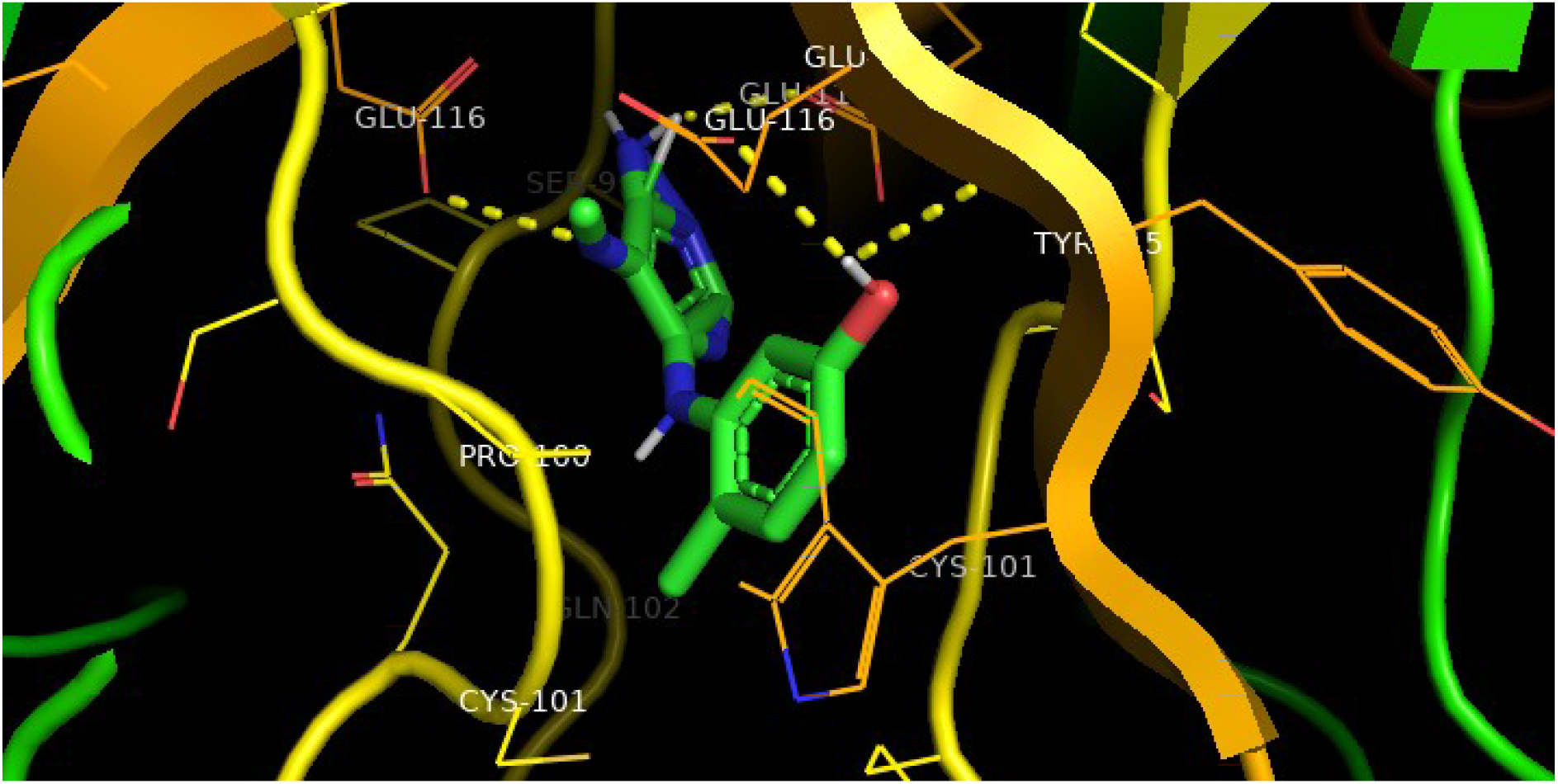
Target-compound interaction of the *de novo* generated SMILES 5-OPT-SMI

**Table 1.** *In Silico* Modelling Target2Drug

| S.No. | Ligand (PubChem CIDs) | Target | Binding Energy (Kcal/mol) |
| --- | --- | --- | --- |
| 1 | 356283 | 1a8m | -8.4 |
| 2 | 11547797 |  | -8.1 |
| 3 | 70681601 |  | -8.6 |
| 4 | 129780132 |  | -8.3 |
| 5 | 144048489 |  | -8.4 |

**Table 2.**
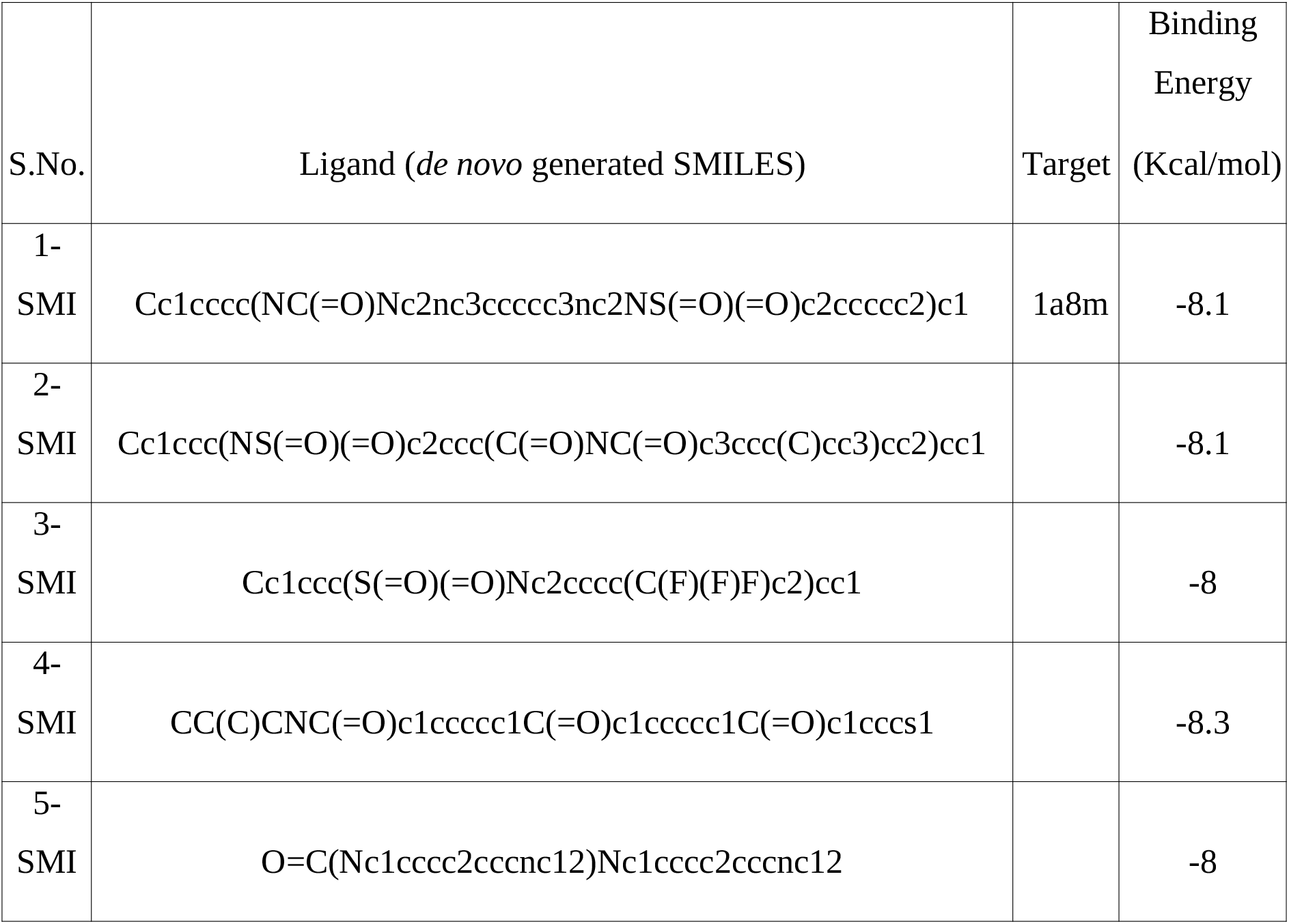
*In Silico* Modelling Target2DeNovoDrug

**Table 2.**
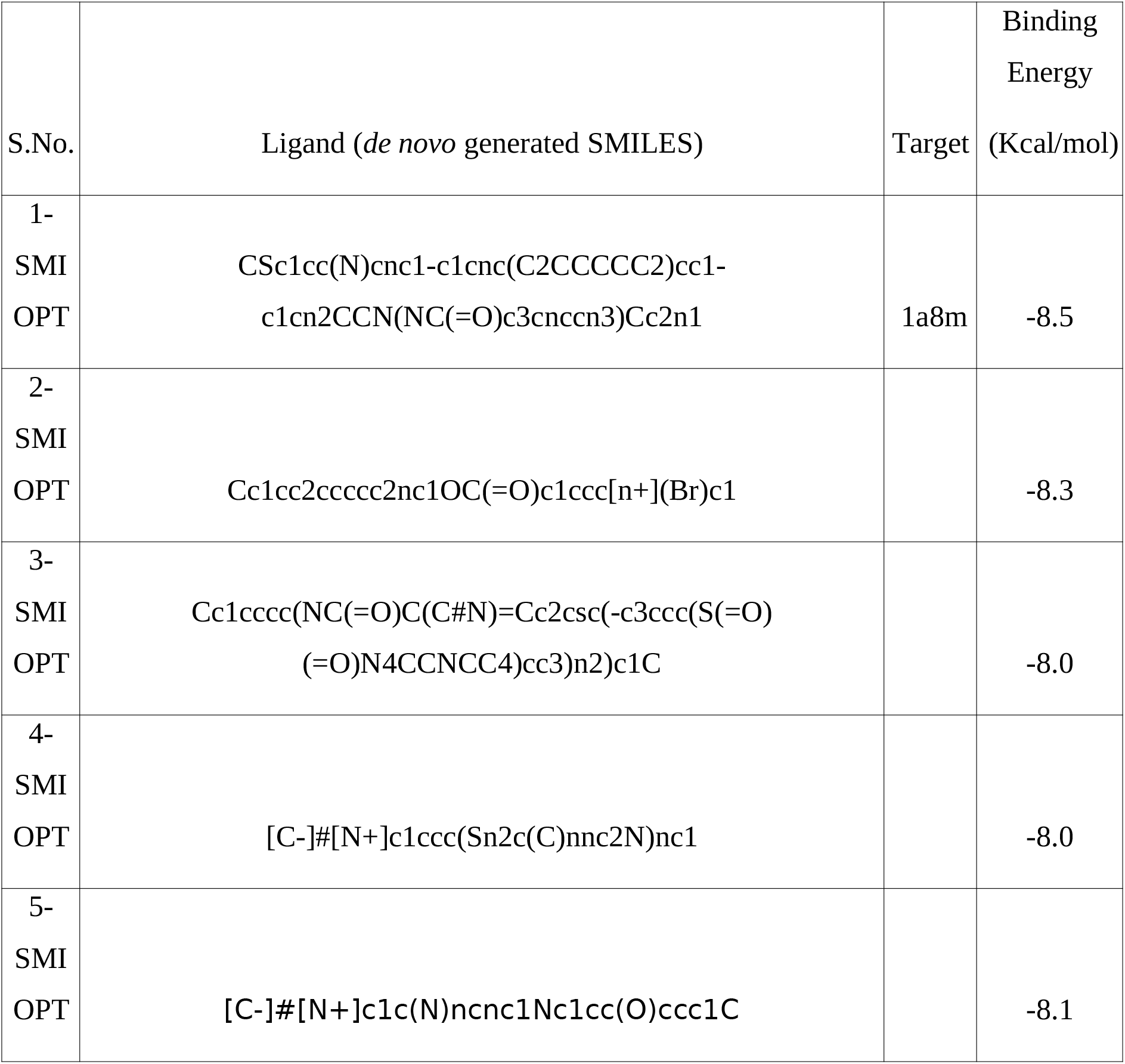
*In Silico* Modelling Target2DeNovoDrugPropMax

The RMSD plot generated from the automated MD protocol of the ‘runGromacs.sh’ bash script of our tool revealing the stability of complex formation involving the protein-ligand complex associated with the complex with the lowest binding energy value from the virtual screening is presented below in Fig.5a and Fig.5b.

**Fig.5a.**
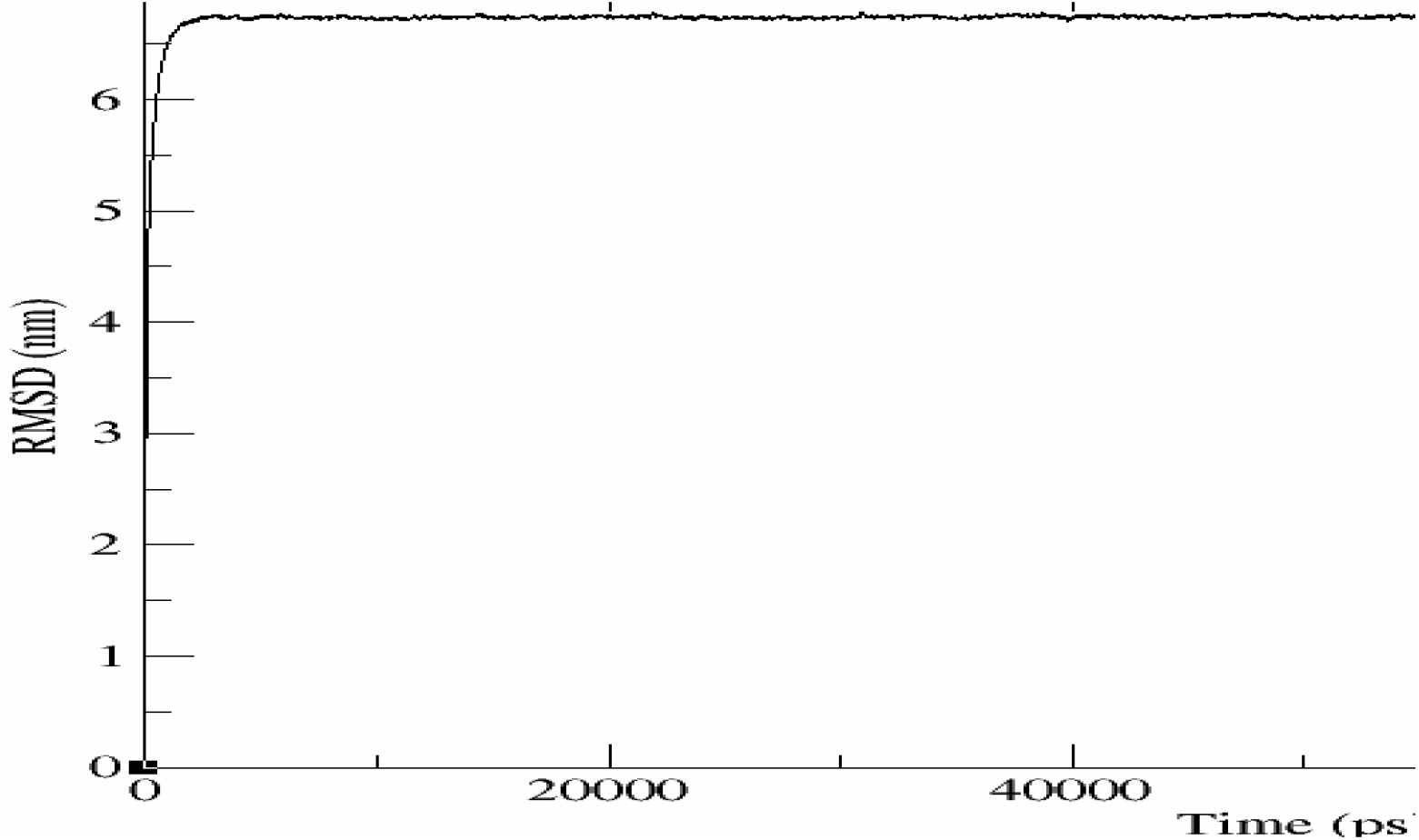
RMSD stabilization of protein-ligand complex associated with the complex of lowest binding energy value

**Fig.5b.**
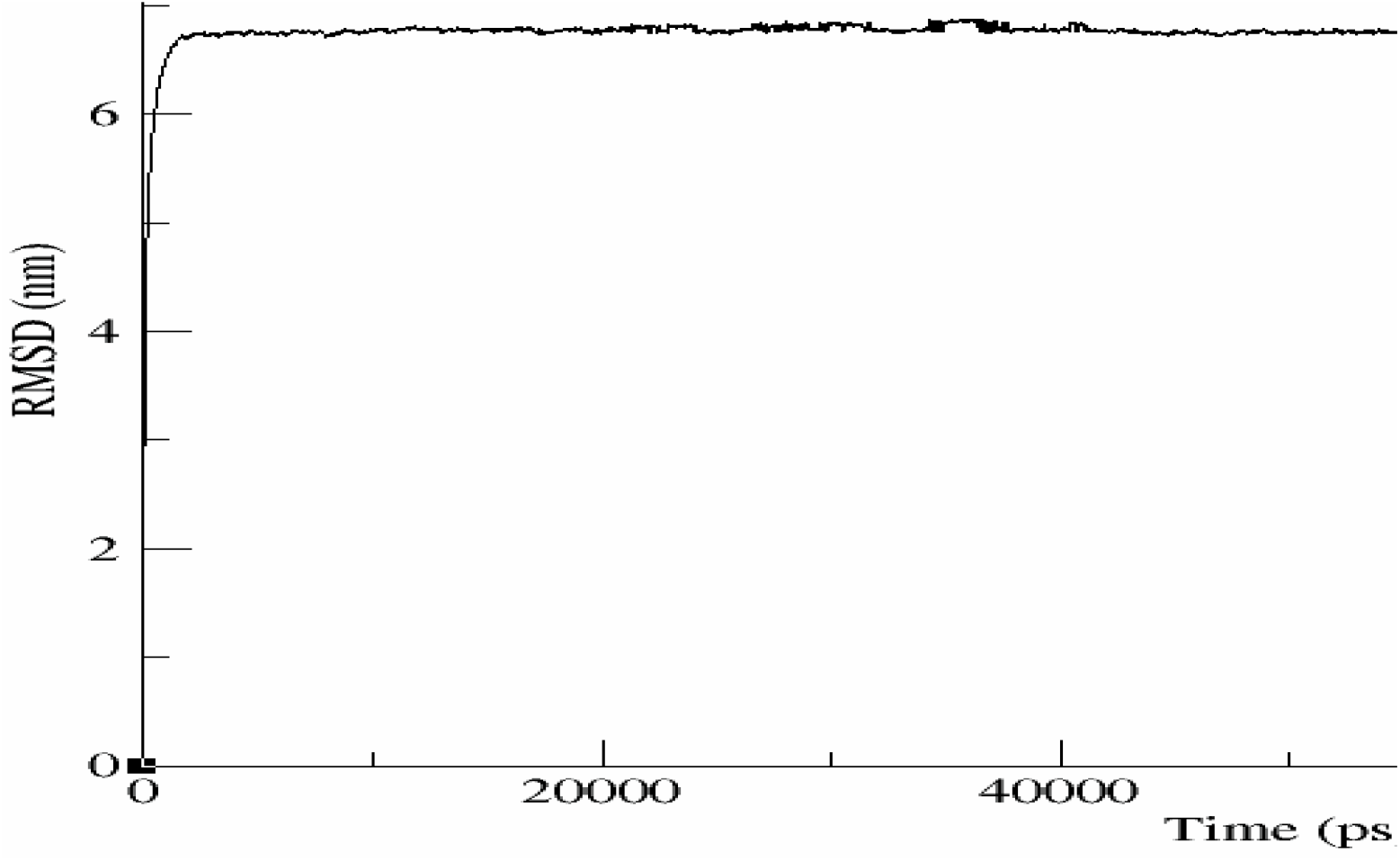
RMSD stabilization of protein-ligand complex associated with the complex of lowest binding energy value

Thus a demonstrated use of the tool *in silico*-deep learning based *de novo* drug design for a target has been shown.

## Conclusion and future outlook

The past decade has seen a surge in the range of application data science, machine learning, deep learning and AI methods to drug discovery. This greatly enhances the tool set already in offer from the computational era of drug discovery such as in silico modeling and therefore a tool which incorporates the best of what both worlds of computational modeling and AI can offer to drug discovery can greatly help researchers in identifying new drug candidates. The presented work involves a assemblage of a variety of AI methods for drug discovery along with an incorporation of in silico techniques to provide a holistic tool for automated drug discovery. A demonstrated use of the tool has been shown with the target signatures of Tumor Necrosis Factor-Alpha, an important therapeutic target in the case of anti-inflammatory treatment. The future scope of the tool involves, running the tool on a High Performance Cluster for all known target signatures to generate data that will be useful to drive AI and Big data driven drug discovery

